# Novel virus-like particle vaccine encoding the circumsporozoite protein of *Plasmodium falciparum* is immunogenic and induces functional antibody responses

**DOI:** 10.1101/2020.12.14.422617

**Authors:** Liriye Kurtovic, David Wetzel, Linda Reiling, Damien R. Drew, Catherine Palmer, Betty Kouskousis, Eric Hanssen, Bruce D. Wines, P. Mark Hogarth, Manfred Suckow, Volker Jenzelewski, Michael Piontek, Jo-Anne Chan, James G. Beeson

## Abstract

RTS,S is the leading malaria vaccine in development, but has demonstrated only moderate protective efficacy in clinical trials. RTS,S is a virus-like particle (VLP) that uses the human hepatitis B virus as scaffold to display the malaria sporozoite antigen, circumsporozoite protein (CSP). Particle formation requires fourfold excess scaffold antigen, and as a result, CSP represents only a small portion of the final vaccine construct. Alternative VLP or nanoparticle platforms that reduce the amount of scaffold antigen and increase the amount of the target CSP antigen present in particles may enhance vaccine immunogenicity and efficacy. Here, we describe the production and characterization of a novel VLP that uses the small surface antigen (dS) of duck hepatitis B virus to display CSP. The CSP-dS fusion protein successfully formed VLPs without the need for excess scaffold antigen, and thus CSP represented a larger portion of the vaccine construct. Importantly, this is the first report of a dS-based vaccine that formed particles without excess scaffold protein. CSP-dS formed large particles approximately 31-74 nm in size and were confirmed to display CSP on the surface. The CSP-dS VLP was highly immunogenic in mice and induced antibodies to multiple regions of CSP, even when administered at a lower vaccine dosage. Vaccine-induced antibodies demonstrated functional activity, including the ability to interact with complement and Fcγ-receptors, both previously identified as important in malaria immunity. Our novel platform to produce VLPs without excess scaffold protein has wide implications for the future development of innovative vaccines for malaria and other infectious diseases.

## INTRODUCTION

There were an estimated 228 million cases of malaria and 405,000 deaths in 2018, largely attributed to infection with *Plasmodium falciparum* (1). Various antimalarial interventions are currently available, including vector control measures to prevent transmission of the malaria-causing parasite from mosquitoes to humans, and drugs to clear infection and treat clinical illness. However, the effectiveness of these tools are threatened by the emergence and spread of drug and insecticide resistance (2), and disruptions to and gaps in public health interventions (3). A critical step towards reducing malaria burden and achieving global malaria elimination will require innovative research to develop new interventions, including efficacious malaria vaccines (4).

The most advanced malaria vaccine in development is RTS,S, which is based on the major surface antigen expressed by *P. falciparum* sporozoites, the circumsporozoite protein (CSP). This vaccine approach targets the initial asymptomatic stage of infection in the liver and therefore aims to prevent parasite replication in the blood and subsequent onset of clinical disease (5). The RTS,S vaccine construct is a fusion protein of a domain derived from the CSP (including the central-repeat and C-terminal regions of the protein) and the human hepatitis B surface antigen (HBsAg), which is co-expressed with excess HBsAg to self-assemble into virus-like particles (VLPs) (6). These particles are ~20 nm in size and co-administered with the potent AS01 adjuvant to enhance vaccine immunogenicity (7). RTS,S/AS01 is the only malaria vaccine to complete phase III clinical trials, which were conducted at 11 study sites in sub-Saharan Africa. RTS,S vaccination conferred ~30-50% protective efficacy against clinical malaria in young children and infants depending on the age group and duration of follow-up (8). However, vaccine efficacy waned substantially in the 12 months after completion of primary vaccination of three doses (9). To further evaluate vaccine safety and efficacy, pilot implementation of RTS,S recently commenced in Ghana, Kenya and Malawi as recommended by the World Health Organization (WHO) (10).

While RTS,S is the leading malaria vaccine candidate, it is an imperfect vaccine that is unlikely to drive malaria elimination as a stand-alone tool. Strategies to modify RTS,S could lead to the development of RTS,S-like or next-generation vaccines with improved immunogenicity, efficacy and longevity in target populations. An important limitation of the RTS,S vaccine construct is the need for excess HBsAg for particle formation to occur and thus, the resulting VLP is comprised of only ~20% CSP-HBsAg fusion protein and ~80% unfused HBsAg scaffold protein (11). A novel CSP-based particle vaccine was recently described known as R21, which did not require excess unfused HBsAg for particle formation, and therefore had an increased proportion of CSP compared to RTS,S. R21 was immunogenic and efficacious in a pre-clinical murine vaccine study, and may improve upon current vaccine approaches (12). Another consideration is the VLP scaffold protein used to display the CSP antigen, which can influence particle size and structure, and therefore influence the induction of protective immune responses (13).

Here we describe the production and characterization of a novel VLP-based vaccine candidate that displays CSP on the surface (**Figure 1**). We used the membrane integral small surface protein (dS) of the duck hepatitis B virus (DHBV) as a VLP scaffold protein (14–16). As a proof-of-concept for this vaccine platform, the same portion of CSP that is included in RTS,S (central-repeat and C-terminal regions of the protein) was selected as the malaria vaccine antigen and was genetically fused to the dS. The resulting CSP-dS fusion protein was overexpressed in recombinant yeast *Hansenula polymorpha*, which is a cell line suitable for large-scale vaccine manufacturing. This approach resulted in the formation of VLPs with the CSP-dS fusion protein alone, without the need for excess dS as a scaffold protein, thereby achieving a higher ratio of the CSP vaccine antigen compared to the approach used for RTS,S and other VLP-based vaccines. The biophysical properties of these VLPs were characterized and the expression of CSP confirmed through Western blotting, transmission electron microscopy and super resolution microscopy. We performed an immunogenicity study of the CSP-dS VLPs in mice and measured the ability of vaccine-induced antibodies to mediate effector functions that have been associated with RTS,S vaccine efficacy (17), such as interacting with complement and Fcγ-receptors (FcγRs) that are expressed on immune cells.

**Figure 1.**
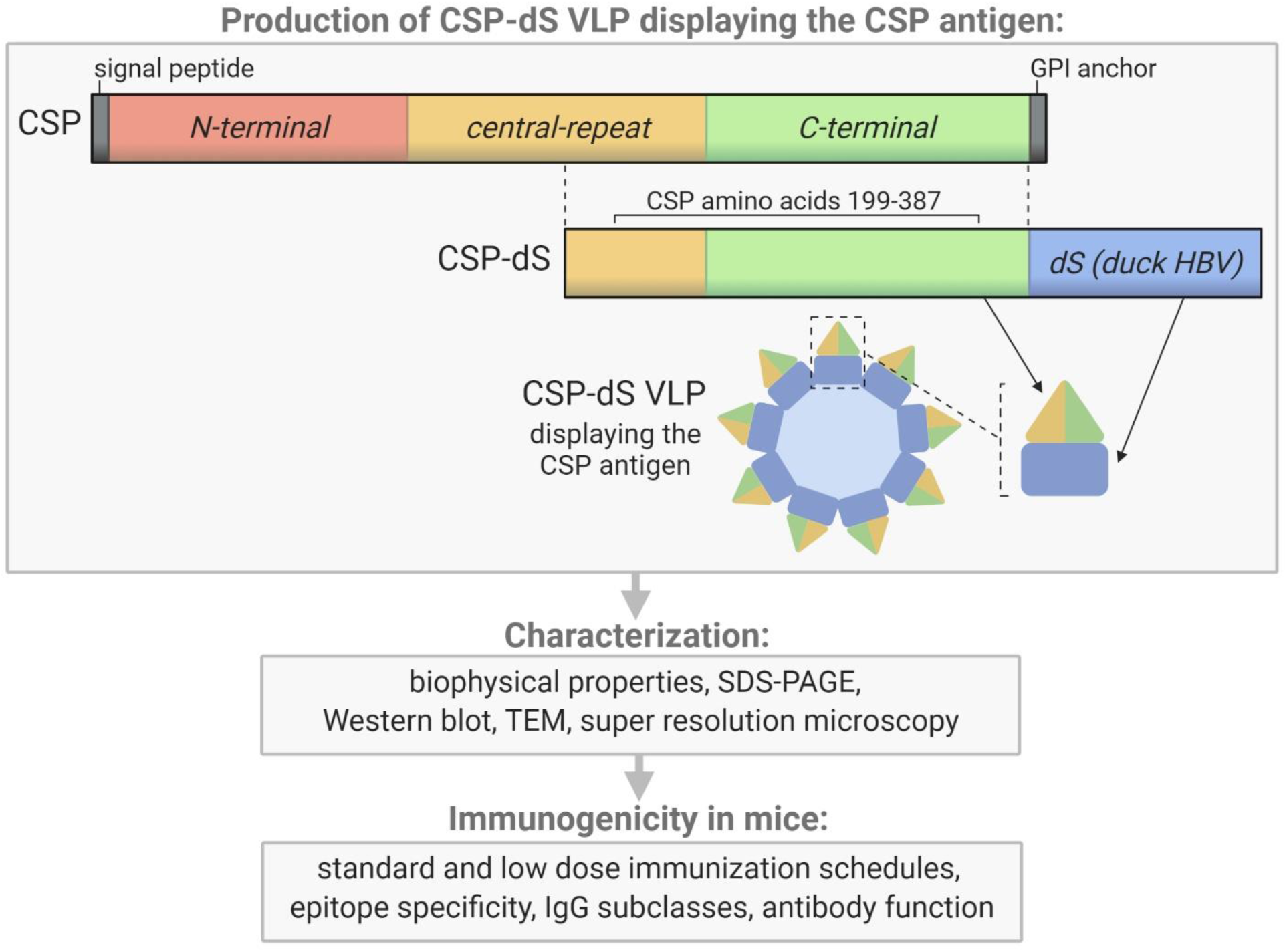
Schematic of the CSP-dS VLP and summary of the study workflow. CSP-dS VLPs were produced by genetically fusing the CSP (including the central-repeat and C-terminal regions, amino acids 199-387) to the small surface protein (dS) of the duck hepatitis B virus (HBV). The CSP-dS fusion protein formed virus-like particles, which were characterized and tested for immunogenicity in mice. Created with BioRender.com.

## MATERIALS AND METHODS

### Generation and purification of the CSP-dS VLP

The CSP-dS fusion protein was composed of amino acids 199-387 of *P. falciparum* CSP (XP_001351122.1) (18) N-terminally fused to the scaffold protein dS (Genbank accession number: MF510122). The resulting CSP-dS encoding gene (Genbank accession number: MH142263) was synthesized by GeneArt/Life Technologies (Regensburg, Germany) and codon-optimized for heterologous expression in *H. polymorpha*. The sequence was subcloned into a derivative of the *H. polymorpha* expression plasmid pFPMT121 (19) which carried the *LEU2* instead *URA3* gene for selection in yeast. The auxotrophic *H. polymorpha* strain ALU3 (relevant genotype: *ade1*, *leu2*, *ura3*) (20) derived from wild type strain ATCC^®^ 34438™ (CBS 4732, IFO 1476, JCM 3621, NBRC 1476, NCYC 1457, NRRL Y-5445) was used as an expression host. Recombinant yeast cell lines were generated by electroporation (21) and a subsequent strain generation and isolation protocol (22). Thereby, the expression plasmid stably integrated into the yeast genome. Heterologous yeast strains were stored as glycerol stocks at − 80 °C.

The production cell line was cultured in 2 L baffled shake flasks filled with 200 mL animal component-free YPG medium containing 20 g L^−1^ glycerol (AppliChem, Germany) as a carbon source and 0.1 g L^−1^ adenine (AppliChem). Pre-cultures grown in YPD medium to stationary phase were used as inoculum. Cultures were incubated at 37 °C and 130 rpm with 5 cm throw. After 56 h of batch growth and derepression of the promoter system by consumption of glycerol, 1% (v/v) methanol was added to the cultures for induction of target gene expression. After 72 h total cultivation time, cells were harvested by centrifugation (6,000*g*, 15 min, 4 °C), washed once with wash buffer (50 mM Na-phosphate buffer, 2 mM EDTA, pH 8.0) and stored at −20 °C.

CSP-dS VLPs were isolated by ultracentrifugation as described previously (15). Briefly, cells were disrupted by high-pressure homogenization. The soluble material was then layered on top of sucrose cushions (2 mL 70% (w/v); 3 mL 20% (w/v)) and the boundary layers between the two sucrose layers were harvested after ultracentrifugation (90 min, 51,000 rpm, 18 °C, Optima™ L90K centrifuge, rotor type: 70.1 Ti, tubes: 16 * 76 mm, Beckman Coulter, USA). These fractions were subsequently mixed with 6 M CsCl (AppliChem) stock solution to 1.5 M final CsCl concentration. Mixtures were subjected to density gradient separation (65 h at 48,400 rpm, 4 °C). Product containing fractions were pooled and desalted by dialysis (Slide-A-Lyzer™ dialysis cassettes, MWCO 20 kDa, Thermo Fisher Scientific, USA) against desalting buffer (8 mM Na-phosphate buffer pH 7, 154 mM NaCl, AppliChem) and sterile filtered (Filtropur S 0.2 filters, Sarstedt, Germany).

### Visualization of CSP-dS VLPs

CSP-dS VLPs were visualized by negative staining transmission electron microscopy (TEM) and super-resolution microscopy (Structured-Illumination Microscopy; SIM) as previously described (14, 15). N-SIM was used to evaluate the co-localization of CSP and the scaffold protein dS in nano-scaled structures, as previously described (15). Samples were dual labelled with 4 μg/mL of primary antibodies (rabbit anti-CSP and mouse anti-dS, 7C12), followed by secondary antibodies (1/1000, anti-rabbit AlexaFluor 594 and anti-mouse AlexaFluor 488; Thermo Fisher Scientific). The super-resolution images were collected using a Nikon N-SIM microscope equipped with 488, 561 and 640 nm lasers, an Andor iXON DU897 EM-CCD camera and a 100x oil immersion lens having a numerical aperture of 1.49. The z-series was acquired using NIS-Elements and analyzed both using NIS-Elements and the open java source, ImageJ/FIJI.

### Western blots

CSP-dS VLPs and monomeric recombinant CSP (rCSP) were prepared under reducing conditions, separated by SDS-PAGE using 4-12% Bis-Tris gels (NuPAGE, Thermo Fisher Scientific) and transferred onto nitrocellulose membranes using the iBlot system (Thermo Fisher Scientific) according to the manufacturer instructions. Membranes were blocked with 10% (w/v) skim milk in phosphate buffered saline (PBS) and probed with polyclonal rabbit anti-CSP IgG (1 μg/mL) and a dS-specific mouse monoclonal antibody (7C12; 1/1000). This was followed by species-specific detection HRP-conjugated antibodies (1/5000; Millipore, USA). Protein bands were detected using SuperSignal Chemiluminescent HRP substrate (Thermo Fisher Scientific) and imaged using the ChemiDoc System (Bio-Rad, USA).

### Evaluation of particle size

Particle size distribution of the CSP-dS VLP preparation was analyzed by dynamic light scattering (DLS) as previously described (15). This analysis was performed before and after analytical high-performance size exclusion chromatography (HP-SEC) using Shimadzu CBM-20A system (Canby, OR, USA) equipped with communication bus modelus (CBM-20A), UV/VIS detector (SPD20-A), degasser (DGU-20A5), liquid chromotograph (LC-20AT), auto sampler (SIL-20AHT) and column oven (CTO-20AC). A volume of 50 μL of the CSP-dS VLP preparation was applied to TSKgel® G6000PWXL 7.8*300 mm column (Tosoh Bioscience GmbH, Griesheim, Germany) in loading buffer (1.7 mM KH2PO4, 7.9 mM Na2HPO4, 2.7 mM KCl; 144 mM NaCl, pH 7.3) at 0.5 ml/min flow. Absorption of the eluate was tracked with the detector SPD20-A at 214 nm and 280 nm wavelength.

### Animal immunizations

Mice were immunized with the CSP-dS VLP or monomeric full-length CSP as a non-VLP control (monomer vaccine) (23). For the first study, Swiss mice received three 10 μg doses of CSP-dS (n=5) or CSP monomer (n=5) vaccines with each dose administered 2 weeks apart (via intraperitoneal injection). The terminal bleed was performed 2 weeks after the final dose and the serum was used for immunogenicity studies. The second study was similar except that C57/BL6 mice were used (due to availability) and instead received three 2 μg doses of CSP-dS (n=5). For all immunizations, the CSP-dS and monomer vaccines were formulated with equal volumes of sterile aluminum hydroxide (Alhydrogel adjuvant; Brenntag, Denmark) and incubated on the shaker for 5 min prior to immunizations. Animal immunizations were conducted at the Animal Facility at the Walter and Eliza Hall Institute (Melbourne, Australia) and ethics approval was obtained by the Animal Ethics Committee of the Walter and Eliza Hall Institute.

### Antigens

The following antigens used in this study were all based on the *P. falciparum* 3D7 sequence: recombinant full-length CSP (excluding the signal peptide and glycosylphosphatidylinositol sequences) expressed in *Escherichia coli* (Gennova, India) (23), synthetic peptide representative of the central-repeat region of CSP (NANP, Life Tein, USA) (24), and recombinant C-terminal region of CSP expressed in HEK293 cells (CT) (24).

### Immunogenicity assays

Antibody responses were measured by standard enzyme-linked immunosorbent assay (ELISA) as follows (24). Ninety-six well flat bottom MaxiSorp Nunc plates (Thermo Fisher Scientific) were coated with 0.5 μg/mL antigen (full-length CSP, NANP and CT) in PBS overnight at 4 °C. Plates were blocked with 1% (w/v) casein in PBS for 2 h at 37 °C and then incubated with test mouse serum diluted in buffer (0.1% casein in PBS) for 2 h at room temperature (RT). To measure total IgG, plates were incubated with goat anti-mouse IgG HRP (Millipore) at 1/2000 in buffer for 1 h at room temperature (RT). Finally, plates were incubated with 2,2’-azino-bis(3-ethylbenzothiazoline-6-sulphonic acid) substrate (ABTS, Thermo Fisher Scientific) for 15 min at RT shielded from light and absorbance was measured at optical density (OD) 405 nm using the Multiskan Go plate reader (Thermo Fisher Scientific). To measure murine IgG subclasses, plates were incubated with goat anti-mouse IgG1, IgG2a, IgG2b or IgG3 detection antibodies (SouthernBiotech, USA) at 1/1000 in buffer, followed by rabbit anti-goat IgG HRP (Sigma-Aldrich, USA) at 1/1000 in buffer, each incubated for 1 h at RT. Finally, plates were incubated with 3,3’,5,5’-Tetramethylbenzidine substrate (TMB, Thermo Fisher Scientific) for 5 min at RT shielded from light; reactivity was stopped using 1 M sulfuric acid and absorbance was measured at OD 450 nm. In all plate-based immunoassays between each incubation step, plates were washed thrice in PBS-Tween20 0.05% (v/v) using the ELx405 automated plate washer (BioTek, USA).

The ability of vaccine-induced antibodies to fix human complement protein, C1q, was performed as previously described (25, 26). Briefly, ninety-six well plates were coated, blocked, and incubated with test serum (at 1/100 dilution) as described for standard ELISA. Following on from this, plates were incubated with 10 μg/mL human C1q (Millipore) in buffer for 30 min at RT. To measure C1q-fixation, plates were then incubated with rabbit anti-C1q IgG (in-house) (24), followed by goat anti-rabbit IgG HRP (Millipore), each at 1/2000 in buffer and incubated for 1 h at RT. Finally, plates were incubated with TMB substrate for 1 h at RT shielded from light; reactivity was stopped using 1 M sulfuric acid and absorbance was measured at OD 450 nm.

Vaccine-induced antibodies were also tested for the ability to interact with human FcγRs, as previously described (17). Briefly, ninety-six well plates were coated, blocked, and incubated with test serum (at 1/100 dilution) as described for standard ELISA. Plates were incubated with 0.2 μg/mL biotin-labelled dimeric recombinant soluble FcγRIIa (H131 allele) (27), and FcγR-binding was detected using streptavidin conjugated HRP (Thermo Fisher Scientific) at 1/10000 dilution for 1 h. Note that for FcγR-biding assays, blocking and dilutions were performed using 1% (w/v) bovine serum albumin in PBS, and all incubations were conducted at 37 °C. Finally, plates were incubated with TMB substrate for 1 h at RT shielded from light; reactivity was stopped using 1 M sulfuric acid, and absorbance was measured at OD 450 nm.

### Statistical analysis

Mouse serum samples were tested in duplicate, and raw data were corrected for background reactivity using no-serum wells that were included as a negative control. Results were shown from two independent experiments, unless specified otherwise. Data were analyzed using GraphPad Prism 8 and antibody responses between mouse vaccine groups were compared using the unpaired t-test where appropriate.

## RESULTS

### Production of the CSP-dS VLP

The recombinant yeast strain, Der#949 (relevant genotype: *ade1*, *LEU2*, *ura3*) was isolated from transformation of host strain ALU3 with the CSP-dS encoding plasmid. This strain was used to produce the CSP-dS VLPs. The CSP-dS construct contained amino acids 199-387 of *P. falciparum* CSP (including the central-repeat and C-terminal regions; **Figure 1**) with the N-terminal MMAP motif and a GPVTN linker between the CSP fragment and the C-terminal dS. Approximately 6 g dry cell weight (DCW) of yeast strain Der#949 were used to isolate 1.2±0.1 mg product VLP containing the CSP-dS fusion protein (Y_P/X_ = ~0.2 mg g^−1^) applying an analytical methodology based on two consecutive steps of ultracentrifugation. The preparation resulted in a clean protein band of approximately 50 kDa in size by SDS-PAGE, indicating there was minimal incorporation of yeast proteins in the VLP preparation as previously shown (15) (**Figure 2A**). Additionally, the CSP-dS fusion protein was specifically recognized by anti-dS and anti-CSP antibodies by Western blot (**Figure 2B**).

**Figure 2.**
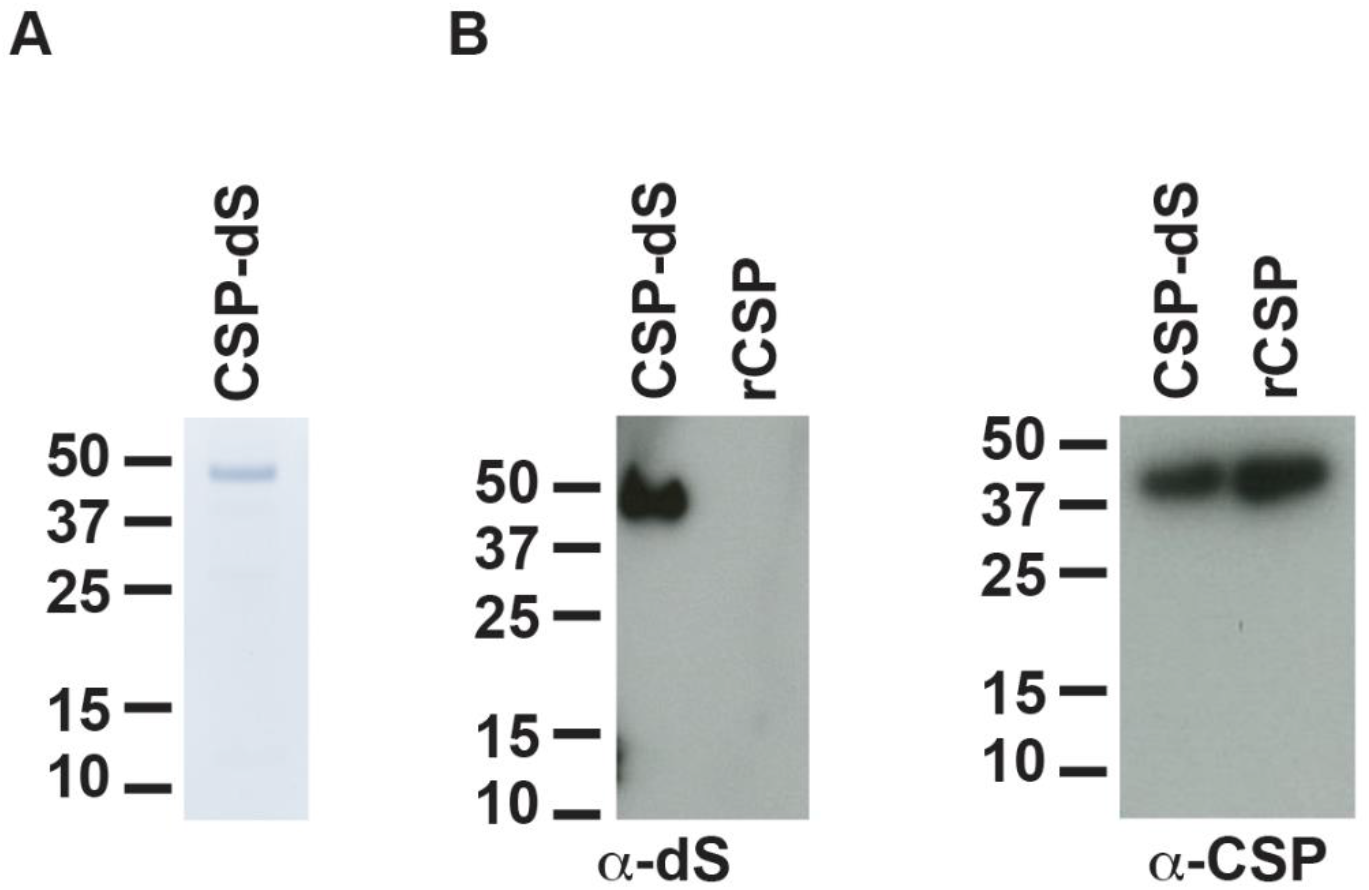
Analyses of purified CSP-dS VLP derived from strain Der#949. (**A**) Reducing SDS-PAGE of CSP-dS VLP purified from strain Der#949 stained with Colloidal blue. A band of expected molecular weight corresponding to CSP-dS was observed at approximately 50 kDa. (**B**) CSP-dS VLP and monomeric recombinant CSP (rCSP) as a control were prepared for Western blot under reducing conditions. Membranes were probed with a monoclonal anti-dS antibody (7C12) and polyclonal anti-CSP antibody.

### CSP-dS formed particulate nanostructures recognized by CSP-specific antibodies

The size distribution analyzed by dynamic light scattering (DLS) revealed a monomodal but polydisperse population of particles (**Figure 3A**; polydispersity index (PDI): 0.59). The VLP preparation was subsequently analyzed by HP-SEC (**Figure S1**) and the product-containing fraction was again analyzed by DLS. The analysis indicated a homogeneous particle population characterized by 73 nm hydrodynamic diameter (PDI: 0.04). Transmission electron microscopy (TEM) was used to characterize the VLPs formed by the CSP-dS fusion protein (**Figure 3B**). The TEM image showed the formation of homogeneous particles of 31-74 nm in diameter based on manual evaluation (**Figure 3B**). The hydrodynamic diameters determined by DLS were slightly larger than the respective diameters specified by manual evaluation of the TEM images, which has been observed previously with other VLP preparations (14). Nevertheless, all data collected were within the dimensions that could be expected for this type of VLP (15, 28). The determined buoyant density (1.14 - 1.15 g cm^−3^) was also plausible for lipoproteins or VLPs (29). Summary of the production process for CSP-dS can be found in **Table 1**.

**Figure 3.**
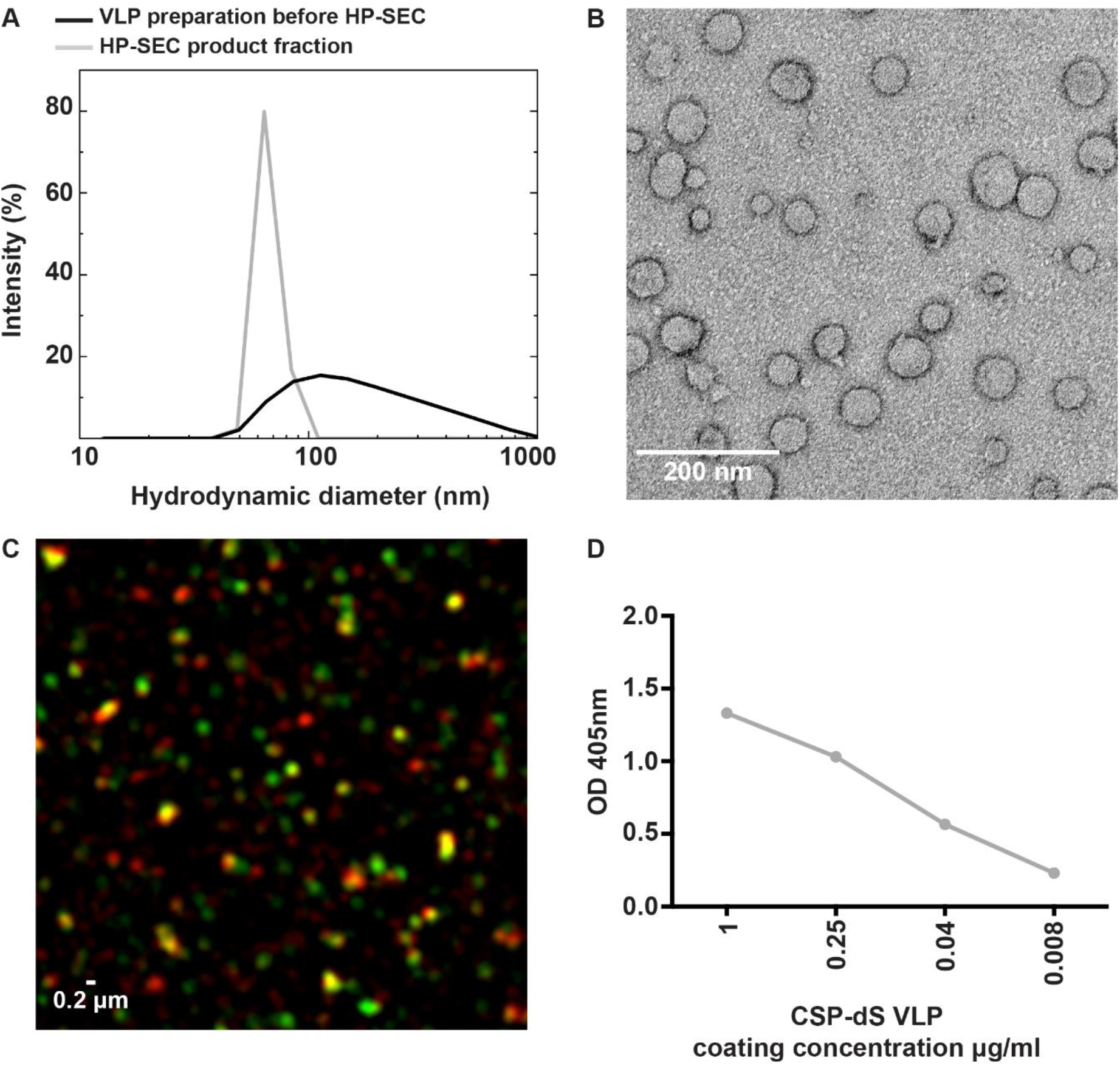
Characterizing the expression of CSP on the VLP surface. (**A**) Analysis of the particulate character of CSP-dS VLP purified from strain Der#949. Size distribution was determined by DLS before and after analysis by HP-SEC. (**B)** The structure of CSP-dS VLPs visualized by negative staining TEM. The VLP structures are consistent with the expected size of approximately 31-74 nm. (**C)** Super-resolution microscopy (N-SIM) was used to visualize CSP-dS VLPs probed with CSP-specific (red) and dS-specific (green) antibodies. Co-localization of CSP and dS are presented in yellow fluorescence. A representative image is shown, and scale bar represents 0.2 μm. (**D**) Binding of polyclonal anti-CSP antibodies to CSP-dS VLP measured by standard ELISA; mean and range of duplicates are shown.

**Table 1.** Summary of production process leading CSP-dS VLP preparation.

| Designation of VLP | CSP-dS VLP |
| --- | --- |
| *H. polymorpha* strain | Der#949 |
| Cell mass generation | Shake flask |
| DCW used for VLP purification [g] | ~6 <sup>(a)</sup> |
| Isolated VLP [mg] | 1.2±0.1 |
| VLP yield per biomass (DCW), $Y_{P/X}$ [mg g<sup>-1</sup>] | ~0.2 <sup>(a)</sup> |
| Product yield per culture volume [mg L<sup>-1</sup>] | ~2 <sup>(a)</sup> |
| VLP diameter by EM [nm] | 31 - 74 |
| Hydrodynamic VLP diameter by DLS [nm] | 73 (PDI: 0.04) <sup>(b)</sup> |
| Buoyant density [g cm<sup>-3</sup>] | 1.14 – 1.15 |

<sup>(a)</sup> Estimated based on determination of OD<sub>600</sub>.
<sup>(b)</sup> Product-containing fraction after analysis by HP-SEC

The structure of the CSP-dS VLPs were further visualized using super-resolution microscopy (N-SIM; **Figure 3C**). Polyclonal anti-CSP antibodies were used to detect CSP expression and monoclonal anti-dS antibodies were used to detect dS expression in the fusion protein. Co-localization of CSP and dS signals was observed in nano-scaled particles, further supporting the formation of VLP structures. We also used polyclonal anti-CSP antibodies to confirm the expression of CSP on the surface of CSP-dS VLPs by standard ELISA (**Figure 3D**).

### CSP-dS VLP was immunogenic and induced antibodies to multiple regions of CSP

Mice were initially immunized with three 10 μg doses of CSP-dS VLP (n=5) or monomeric recombinant CSP (n=5) as a non-VLP comparison control, each formulated with Alhydrogel adjuvant. Sera were collected after the final dose and evaluated for antibodies by standard ELISA. Serum samples were tested between 1/100 and 1/64000 dilution for total IgG to full-length CSP (**Figure 4A**). There was a strong induction of anti-CSP IgG in all mice, apart from mouse #51 of the monomer vaccine group (**Figure S2A**). We next measured the IgG response to antigens representing the central-repeat (NANP) and C-terminal (CT) regions of CSP (**Figure 4B-C**). All mice demonstrated high levels of IgG to the NANP and CT antigens, apart from mouse #51, and mouse #48 of the monomer vaccine group also had low reactivity to the CT (**Figure S2B-C**). Total IgG responses to full-length CSP and NANP were comparable between groups (p=0.671 and p=0.459, respectively), but anti-CT IgG was significantly higher for the CSP-dS vaccine group compared to the monomer vaccine group (p=0.034). We also observed these trends when testing sera collected after only two vaccine doses (**Figure S3**). Antibodies to full-length CSP were further characterized for IgG subclasses (**Figure 5A**). Immunization with the CSP-dS VLP or monomer predominantly induced anti-CSP IgG1, whereas IgG2a, IgG2b and IgG3 responses were variable, which is typical for vaccinations using Alhydrogel as the adjuvant. Overall, there were no significant differences in IgG subclass response between the vaccine groups (p>0.05 for all tests).

**Figure 4.**
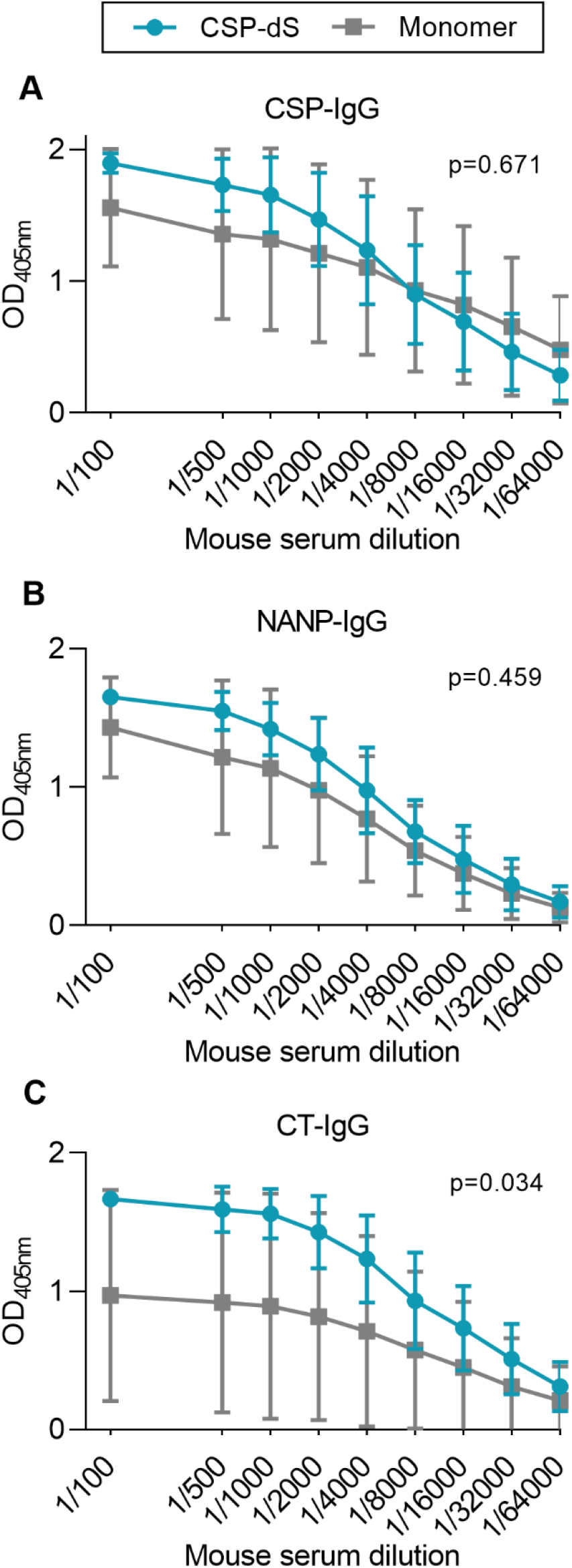
CSP-dS is immunogenic in mice. Swiss mice were immunized with three 10 μg doses of CSP-dS (n=5; circles) or monomeric CSP as a control (n=5; squares). Serum samples collected after the final immunization were tested for total IgG to (**A**) full-length CSP, and antigens representing the (**B**) central-repeat (NANP) and (**C**) C-terminal (CT) regions of CSP. The x-axis is presented on a log2 scale and the mean and standard deviation of two independent experiments are shown. Reactivity between the CSP-dS and monomer vaccine groups were compared using the unpaired t-test (with Welch’s correction).

**Figure 5.**
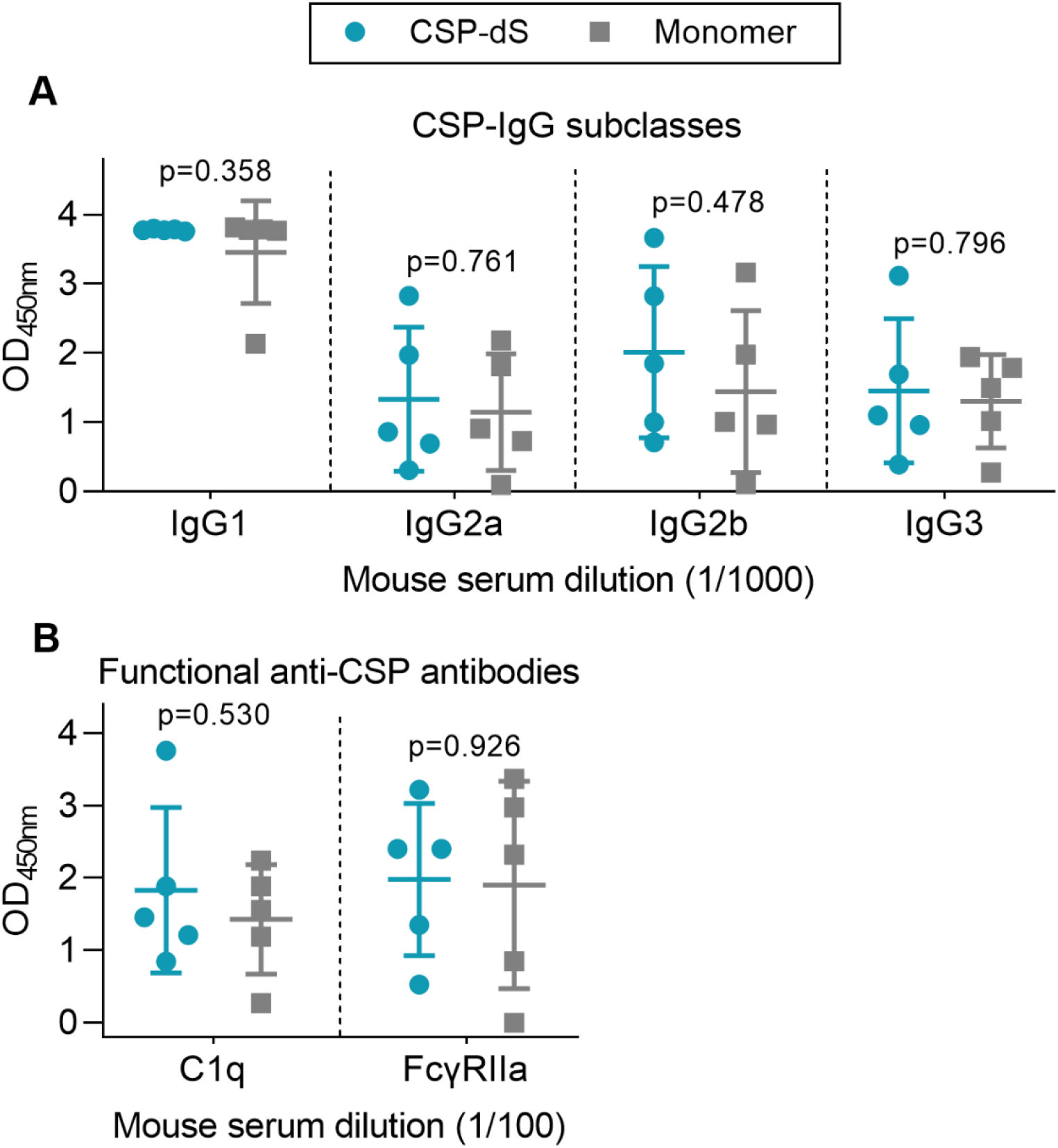
Immunization with CSP-dS induces different IgG subclasses and functional antibody responses. Swiss mice were immunized with three 10 μg doses of CSP-dS (n=5; circles) or monomeric CSP as a control (n=5; squares). Serum samples collected after the final immunization were tested for (**A**) anti-CSP IgG subclass responses and (**B**) functional activity against full-length CSP, including C1q-fixation and binding to dimeric FcγRIIa (data from one experiment). Mean and standard deviation are shown and reactivity between the CSP-dS and monomer vaccine groups were compared using the unpaired t-test.

### Vaccine-induced antibodies had Fc-dependent effector functions

We next examined whether vaccine-induced antibodies could mediate Fc-dependent effector functions, which have been correlated with RTS,S vaccine efficacy (17), using established plate-based detection assays (**Figure 5B**). Firstly, we measured the ability of anti-CSP antibodies to fix human complement protein, C1q. C1q-fixation is essential to activate the classical complement pathway, and has been previously identified as a mechanism of naturally-acquired and vaccine-induced immunity to *P. falciparum* sporozoites (25, 30). The ability of vaccine-induced antibodies to fix C1q was moderate and did not significantly differ between vaccine groups (p=0.530, **Figure 5B**). We then measured whether the antibodies could effectively form immune complexes to CSP that interact with dimeric recombinant human FcγRIIa, as a functional surrogate for antibody cross-linking of FcγR expressed on immune cells. FcγRIIa is widely expressed on different immune cells, such as monocytes and neutrophils, and can interact with antibodies to promote opsonic phagocytosis. Importantly, FcγR-binding and opsonic phagocytosis are known functional mechanisms of antibodies to the CSP antigen (17). There was a moderate level of FcγRIIa-binding that was comparable between vaccine groups (p=0.926, **Figure 5B**). However, it should be noted that mouse IgG subclasses do not have same potential to fix human complement and bind human FcγRs as seen with human IgG subclasses induced by malaria vaccines, which are typically predominantly IgG1 and IgG3 (31).

### Fractional dosing of CSP-dS was immunogenic in mice

To determine whether the CSP-dS VLP was immunogenic at a lower dose, we performed a second immunization study. Mice were immunized with three 2 μg doses of CSP-dS, which was a fraction (1/5^th^) of the original dosage administered in the first vaccine study. Serum samples collected after the final immunization were evaluated by standard ELISA, and all mice demonstrated strong IgG responses to full-length CSP, NANP and CT antigens (**Figure 6A-C**). Therefore, CSP-dS was highly immunogenic, even when administered at a lower dose.

**Figure 6.**
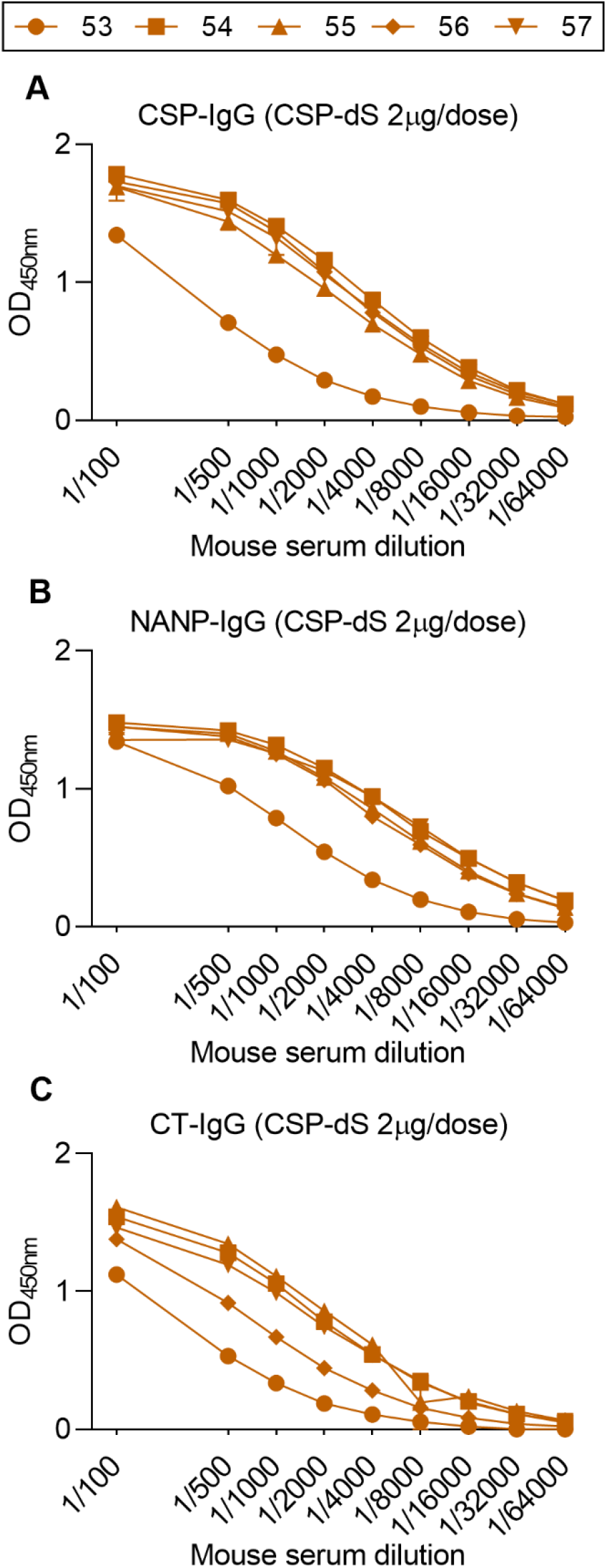
Fractional dosing of CSP-dS is immunogenic in mice. C57/BL6 mice were immunized with three 2 μg doses of CSP-dS (mouse #53-57). Serum samples collected after the final immunization were tested for total IgG to (**A**) full-length CSP, and antigens representing the (**B**) central-repeat (NANP) and (**C**) C-terminal (CT) regions of CSP. The x-axis is presented on a log2 scale and the mean and range of two independent experiments are shown.

## DISCUSSION

New vaccine platforms could facilitate the development and licensure of highly efficacious malaria vaccines. Here, we describe a novel technique to produce VLPs using the small surface antigen (dS) of duck hepatitis B virus as a scaffold to express malaria antigens for vaccine development. As a proof-of-concept, we used CSP as the model antigen since CSP-based vaccines have shown reproducible protective efficacy in animal models, and CSP forms the basis of leading RTS,S malaria vaccine, which has completed phase III clinical trials. Our CSP-dS fusion protein resulted in particle formation without the need for excess unfused dS, and thus a higher density of CSP was displayed on the VLP surface compared to conventional VLP-based vaccine approaches. The CSP-dS VLP was immunogenic in mice and induced antibody responses to multiple regions of the CSP, even when administered at lower doses. Furthermore, vaccine-induced antibodies demonstrated substantial Fc-dependent functional activities.

We report for the first time, the production of a dS-based VLP that formed particles in the absence of excess unfused scaffold protein (32, 33). Previously, we reported the expression of malaria transmission-blocking vaccine antigens Pfs230 and Pfs25 as VLPs using the duck HBV platform (14, 16). However, in those cases, the VLPs were formed with excess dS protein and, as a result, only a minority of the total protein content was formed by the vaccine antigen. The CSP-dS fusion protein successfully formed particulate structures ranging from 31-74 nm as visualized by TEM. We also confirmed the display of CSP on the CSP-dS VLP, and importantly did not detect any unfused dS or degradation productions by Western blot. While the absence of unfused dS made preparation of the CSP-dS VLP a special case, the resulting dimensions and buoyant density were similar to previous dS-based VLP developments (14, 15). Transfer of the analytical purification process to a scalable downstream process for vaccine production represents an important task for the future and may improve the VLP product yield per biomass (Y_P/X_). Nevertheless, considering the CSP-dS VLP was exclusively formed by the fusion protein, the yield per biomass of ~0.2 mg g^−1^ was remarkable. We additionally generated a prototrophic RB11-based strain, which performed like the Der#949 strain regarding CSP-dS production (data not shown) but would be more suited for future scale-up due to simpler nutrient requirements. The CSP-dS VLP vaccine is likely to be suited to formulation with a range of different adjuvants, or to be co-formulated with other vaccine components, such as other malaria antigens, in multivalent vaccines. However, further studies will be needed to assess specific antigen combinations and formulations.

The CSP-dS VLP was immunogenic in mice and induced antibody responses to full-length CSP and antigens representing the central-repeat and C-terminal regions of the protein. Notably, the CSP-dS construct was based on a truncated form of CSP including the central-repeat and C-terminal regions, similar to the RTS,S vaccine construct. However, antibodies to the N-terminal region of CSP can inhibit sporozoite function *in vitro*, and the passive transfer of N-terminal antibodies can confer protection against malaria in murine model (34–37). Therefore, inclusion of the N-terminal region may be favorable for future CSP-based vaccines, or selected epitopes such as the junctional epitope located between the N-terminal and central-repeat regions (37, 38). Additional modifications could also be made in the central-repeat region, which is comprised of ~37 NANP tandem repeats. Indeed, vaccine constructs with fewer repeats can modulate the immune response to other regions of CSP when given to mice (39). Furthermore, it was recently shown that mice vaccinated with a VLP encoding only 9 NANP repeats could induce potent anti-CSP antibodies that fixed complement (40).

CSP-dS vaccine-induced antibodies demonstrated moderate Fc-dependent functional activity, which was somewhat expected given that antibodies were predominately murine IgG1. While human IgG1 is cytophilic and potently activates complement and promotes opsonic phagocytosis (and other Fc-dependent effector functions), murine IgG1 is strikingly different and has little functional activity (31). The complement fixing and Fcγ-receptor binding activity in this study was likely due to IgG2b and IgG3, which are were induced at lower levels. In this study, we used Alhydrogel as an adjuvant, but alternative adjuvants should be further investigated that may alter the IgG subclass response and possibly enhance antibody functional activity. Studies on the recently described R21 vaccine have compared the use of different adjuvants in mouse immunization studies (12). R21 is another VLP-based vaccine similar to RTS,S, but is generated using a different expression system and does not require excess unfused HBsAg scaffold protein to form particles. R21 administered with Alhydrogel induced the lowest antibody titers in BALB/c mice, while oil-in-water emulsion and saponin-based adjuvants induced higher antibody titers that were comparable (12). Interestingly, only the latter formulation conferred protection against sporozoite challenge. This suggested that differences in protective efficacy were not explained by differences in IgG titer but may have been due to differences in IgG subclass or antibody function, which were not explored.

Mice immunized with CSP-dS VLP or monomer CSP as a non-VLP control had comparable antibody responses to full-length CSP and the central-repeat region. However, antibodies to the C-terminal region were significantly higher in the CSP-dS group, in which 5/5 mice had high levels of anti-CT IgG, while for the monomer group only 3/5 mice had strong reactivity. We anticipated to see higher antibody responses in the CSP-dS group because VLPs are generally considered more immunogenic than monomeric protein as they are larger in size, and therefore optimal for uptake by antigen presenting cells and inducing adaptive immune responses (41). Indeed, the CSP-dS VLP was large in size and formed particles ~3 times greater than related CSP-based vaccines, RTS,S and R21. Although, we must take caution when directly comparing the CSP-dS VLP and monomer vaccines because even though both were administered at 10 μg doses, the amount of CSP present in the CSP-dS VLP would have only been ~5.2 μg (as the fusion protein was comprised of ~52% CSP). Therefore, the CSP-dS VLP may have been superior to the monomer vaccine, given that the amount of CSP administered was lower overall. We performed a second immunization study and found that even at a lower concentration of 2 μg per dose, the CSP-dS VLP strongly induced antibodies to full-length CSP and the central-repeat and C-terminal regions in 5/5 mice. We did not undertake a formal comparison of the immunogenicity of CSP-dS VLPs versus monomeric protein since studies in humans have already reported superior immunogenicity of VLPs and other nanoparticle vaccines. Formal assessment of immunogenicity would be a part of any potential future development work.

The selected adjuvant may have also influenced vaccine-induced responses. Early studies found that RTS,S administered with alum was poorly efficacious against controlled human malaria infection in healthy volunteers (11). However, vaccine efficacy was enhanced when RTS,S was administered with an oil-in-water emulsion of MPL and QS-21, which has since been modified to a liposome-based adjuvant known as AS01 and is the current RTS,S vaccine adjuvant (8). This raises an important question of what the true advantages are of using a VLP-based vaccine instead of recombinant protein, and whether enhanced immunogenicity and efficacy are largely due to the adjuvant itself (5). It is difficult to draw a clear conclusion, as many studies do not include a non-VLP vaccine as a negative control.

There are interesting similarities and differences between our CSP-dS VLP and the related R21 and RTS,S vaccines. Notably, CSP-dS and R21 are both comprised of a fusion protein between CSP and scaffold protein that can form particles without the need for excess unfused scaffold protein (12). This may be advantageous as CSP represents a larger portion of the final vaccine construct in comparison to the RTS,S vaccine that is largely comprised of scaffold protein (11). Another possible advantage of our CSP-dS VLP is that the dS scaffold protein is derived from duck hepatitis B virus rather than human hepatitis B virus (HBV), which is used for R21 and RTS,S. The dS scaffold protein forms larger particles that may be more effective at inducing immune responses (41), and the CSP-dS VLPs were ~3 times larger than the R21 and RTS,S VLPs (12). Furthermore, the dS scaffold may also be favorable as vaccine recipients should not have pre-existing antibodies to the duck HBsAg. In some settings, it may not be desirable to use a malaria vaccine that includes HBsAg in populations who have already received HBV vaccines. HBV vaccines are recommended at birth in many regions, whereas malaria vaccines would be given later in infancy. It is unclear whether pre-existing serum antibodies to the human HBsAg would interfere with vaccine-induced responses, but this has been reported for serum antibodies to CSP (42, 43).

In summary, we have demonstrated a novel approach to develop VLP structures without the co-expression of unfused dS scaffold protein. We produced a CSP-dS fusion protein that successfully formed homogenous particles and correctly displayed CSP on the VLP surface. Furthermore, the CSP-dS VLP was immunogenic in mice and induced potent IgG responses to multiple regions of the CSP antigen, which demonstrated functional antibody responses. Our platform to produce VLPs formed with the fusion protein only, without the need for excess scaffolding protein, is highly novel and warrants further evaluation in pre-clinical efficacy studies for CSP and other candidate malaria vaccine antigens.

## Acknowledgements

The authors gratefully acknowledge Brigitte Derksen for technical assistance and Jack Richards for providing mouse IgG subclass reagents. Recombinant full length CSP was kindly provided by PATH Malaria Vaccine Initiative (USA) and Gennova Biopharmaceuticals (India).

## Author contributions

LK performed immunoassays for both mouse immunization studies and data analysis. DW generated, purified, and characterized the CSP-dS VLP. LR performed immunoassays for the first mouse immunization study. DRD generated recombinant proteins used in immunoassays. CP, BK, EH and JAC visualized the VLPs using N-SIM and TEM. BDW and PMH generated reagents for immunoassays. JAC and DRD performed SDS-PAGE and Western blotting analyses. DW, MS, VJ, MP, JAC and JGB were involved in the study design. LK, DW, JAC and JGB wrote the manuscript, which was reviewed and approved by all authors.

## SUPPLEMENTARY FIGURES

**Figure S1.**
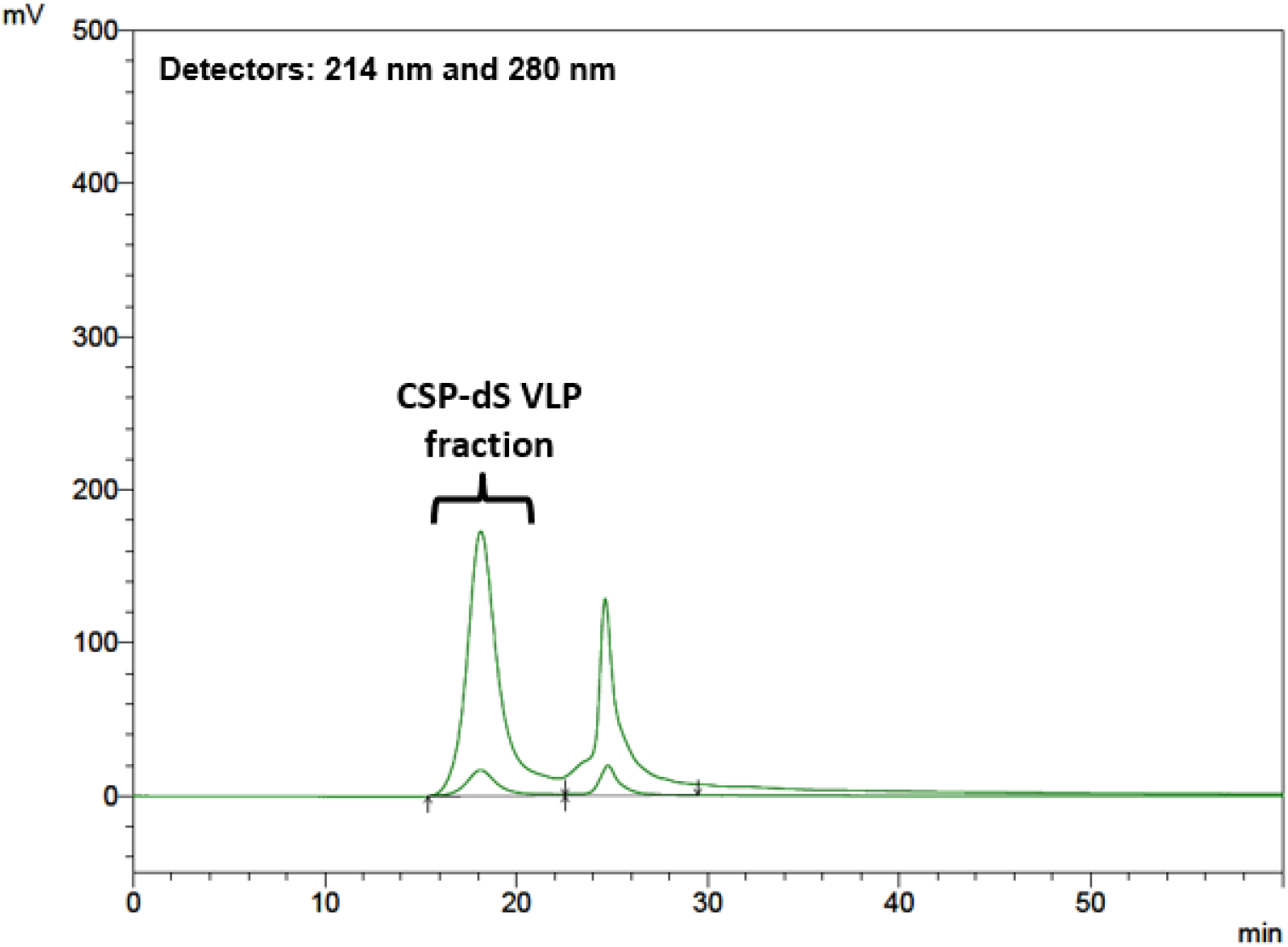
Chromatogram of the HP-SEC analysis of the CSP-dS VLP preparation. Elution profiles were recorded at 214 nm (peptide bonds) and 280 nm (aromatic amino acids). Fractions were analyzed by SDS-PAGE (data not shown) and the marked peak was identified as product-containing fraction and analyzed by DLS.

**Figure S2.**
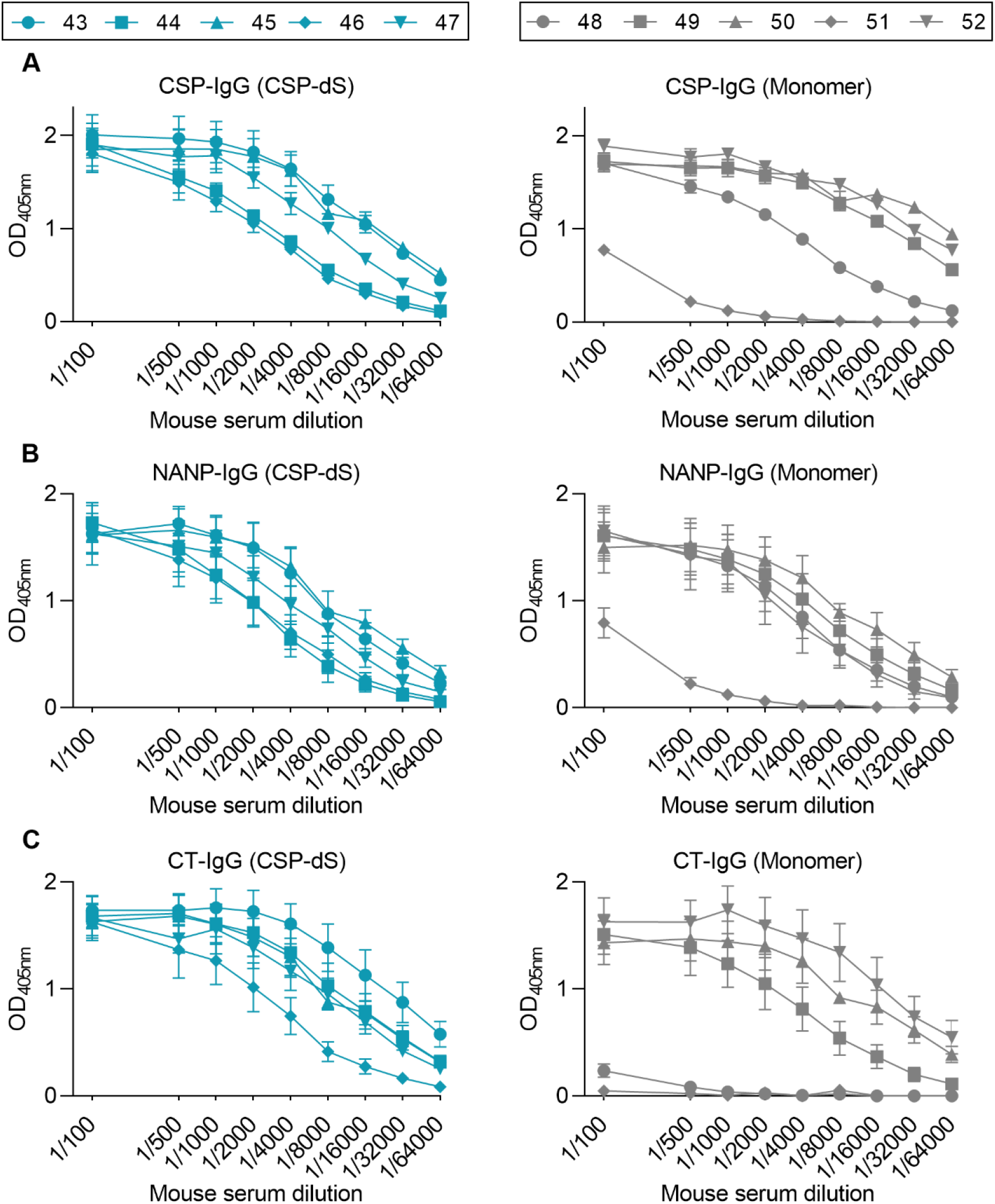
Antibody data from individual mice immunized with CSP-dS and monomer vaccines. Swiss mice were immunized with three 10 μg doses of CSP-dS (mouse #43-47) or monomeric CSP as a control (mouse #48-52). Serum samples collected after the final immunization were tested for total IgG to (**A**) full-length CSP, and antigens representing the (**B**) central-repeat (NANP) and (**C**) C-terminal (CT) regions of CSP. The x-axis is presented on a log2 scale and the mean and range of two independent experiments are shown.

**Figure S3.**
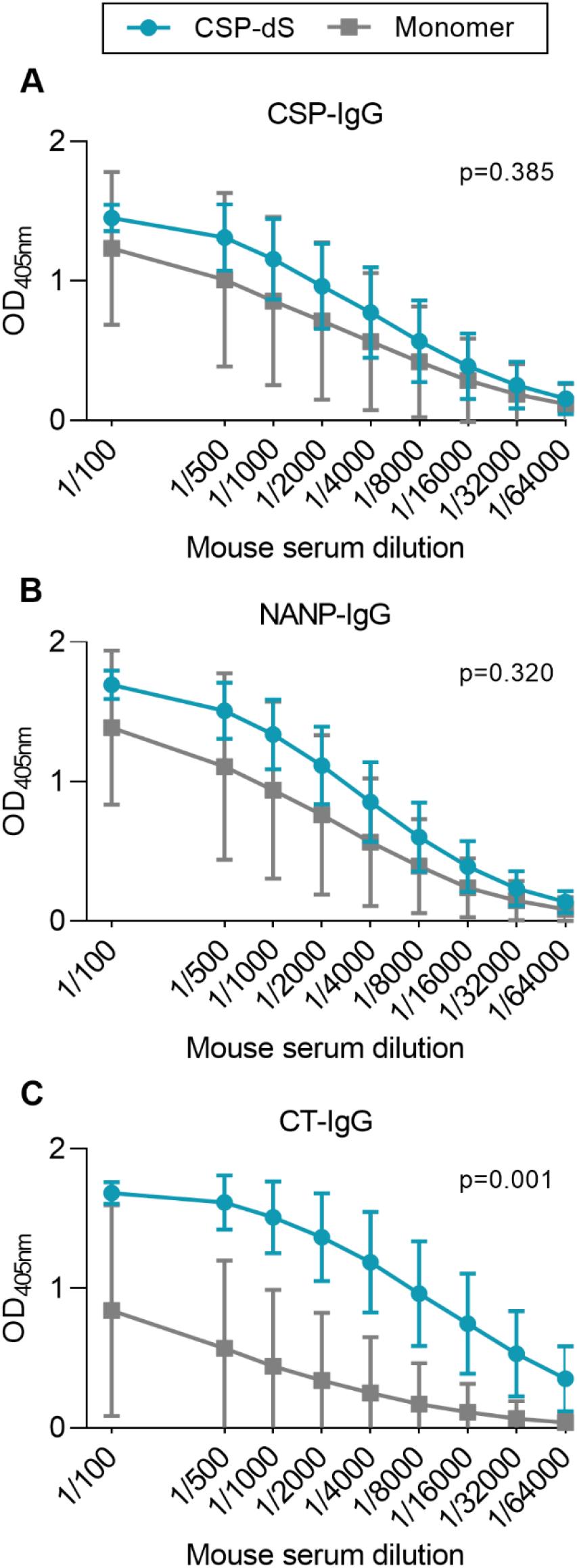
Immunogenicity of CSP-dS after two doses. Swiss mice were immunized with three 10 μg doses of CSP-dS (n=5; circles) or monomeric CSP as a control (n=5; squares). Serum samples collected after the second immunization were tested for total IgG to (**A**) full-length CSP, and antigens representing the (**B**) central-repeat (NANP) and (**C**) C-terminal (CT) regions of CSP. The x-axis is presented on a log2 scale and the mean and standard deviation of one experiment is shown. Reactivity between the CSP-dS and monomer vaccine groups were compared using the unpaired t-test (with Welch’s correction).

